# Defining the toxicity limits on microbial range in a metal-contaminated aquifer Running Title: Inorganic ion toxicity limits on microbial range

**DOI:** 10.1101/388306

**Authors:** Hans K. Carlson, Morgan N. Price, Mark Callaghan, Alex Aaring, Romy Chakraborty, Hualan Liu, Adam P. Arkin, Adam M. Deutschbauer

## Abstract

In extreme environments, toxic compounds restrict which microorganisms persist. However, in complex mixtures of inhibitory compounds, it is challenging to determine which specific compounds cause changes in abundance and prevent some microorganisms from growing. We focused on a contaminated aquifer in Oak Ridge, Tennessee, U.S.A. that has low pH and high concentrations of uranium, nitrate and many other inorganic ions. In the most contaminated wells, the microbial community is enriched in the *Rhodanobacter* genus. *Rhodanobacter* relative abundance is positively correlated with low pH and high concentrations of U, Mn, Al, Cd, Zn, Ni, Co, Ca, NO_3_^−^, Mg, Cl, SO_4_^2−^, Sr, K and Ba and we sought to determine which of these correlated parameters are selective pressures that favor the growth of *Rhodanobacter* over other taxa. Using high-throughput cultivation, we determined that of the ions correlated high *Rhodanobacter* abundance, only low pH and high U, Mn, Al, Cd, Zn, Co and Ni (a) are selectively inhibitory of a sensitive *Pseudomonas* isolate from a background well versus a representative resistant *Rhodanobacter* isolate from a contaminated well, and (b) reach toxic concentrations in the most contaminated wells that can inhibit the sensitive *Pseudomonas* isolate. We prepared mixtures of inorganic ions representative of the most contaminated wells and verified that few other isolates aside from *Rhodanobacter* can tolerate these 8 parameters. These results clarify which toxic inorganic ions are causal factors that impact the microbial community at this field site and are not merely correlated with taxonomic shifts.

## Introduction

Microorganisms survive within an n-dimensional biogeochemical space comprised of diverse nutrients and stressors (Eisenhauer *et al.*, 2012). While neutral effects such as drift, dispersal and speciation can alter microbial community composition, selective pressures are important to identify because they represent the biogeochemical determinants that influence microbial community composition based on variations in the relative fitness of microbial sub-populations (Vellend, 2010). For example, the presence or absence of preferred electron donors, electron acceptors, trace nutrients, toxic compounds and other parameters such as pH and temperature impact the relative success of different microbial sub-populations and thereby alter microbial community composition (Fierer, 2017). Often selective pressures are identified based on correlations between the measurements of the relative abundance of different taxa and measurements of geochemical parameters, but correlative analysis does not identify causal relationships, and many observed correlations can be misleading. Thus, laboratory measurements of microbial fitness in biogeochemical gradients are usually needed to confirm to the causality of selective pressures that influence microbial community composition. Given the complexity of the environmental systems, high-throughput approaches are needed to evaluate the relative fitness of microbial sub-populations in multi-dimensional gradients (Carlson *et al.*, 2017a).

In both pristine and anthropogenically perturbed environments, toxic inorganic ions can inhibit microbial growth, survival and activity and thereby impact the composition of microbial communities (Gadd, 2010; Smith *et al.*, 2015). Many soils and aquifers have naturally high levels of diverse toxic ions, and, depending on the prevailing geochemical conditions, these ions can be solubilized and impact biological systems (Tchounwou *et al.*, 2012; Duxbury, 1985). For example, in the rhizosphere, inorganic ions are solubilized from minerals by organic acids released from plant roots. In this acidic and high metal environment, low pH and metal tolerant taxa are favored (Carson *et al.*, 2009; Hinsinger *et al.*, 2003). Contaminated environments vary even more in pH and inorganic ion concentration, and taxonomic shifts are correlated with changes in ion concentrations in wastewater treatment plants (Stepanauskas *et al.*, 2005), urban estuaries (Duxbury, 1985), acid mine drainage (Baker and Banfield, 2003; Johnson and Hallberg, 2003) and metal contaminated aquifers (Li *et al.*, 2018; Hug *et al.*, 2015; Smith *et al.*, 2015; Lin *et al.*, 2012). Studies into the microbial ecology of these sites is important for understanding and mitigating the impact of ion toxicity on ecosystem health (Atashgahi *et al.*, 2018), but they also represent useful sites for testing general principles of microbial ecology.

The aquifer at the U.S. Department of Energy Field Research Center in Oak Ridge, Tennessee (ORFRC) is both low pH and metal contaminated (Brooks, 2001). At the ORFRC, between 1951 and 1983, nitric acid solubilized uranium waste from the Y-12 nuclear processing plant was deposited in the unlined S-3 ponds along with mixed metal and organic wastes from other DOE facilities (Brooks, 2001). The intrusion of low pH waste further dissolved the fractured shale and karst aquifer matrix releasing other inorganic ions. For example, dissolution of clay minerals releases Mn, Al and other trace metals (Gromet *et al.*, 1984; Reinhard *et al.*, 2013) while dissolution of carbonates releases alkali earth metals such as Ca and Mg (Gromet *et al.*, 1984). Consequently, the aquifer’s pH ranges from 3-11 and the concentration of many inorganic ions in groundwater span over 5 orders of magnitude (Thorgersen *et al.*, 2015; Smith *et al.*, 2015). Identifying which toxic inorganic ions impact the microbial community at this field site is important for interpreting microbial community datasets (Smith *et al.*, 2015; Li *et al.*, 2018), for understanding the selective pressures on metal resistance (Thorgersen *et al.*, 2017; Hemme *et al.*, 2016; 2010; Price *et al.*, 2018) and for designing optimum bioremediation strategies (Williams *et al.*, 2013).

A number of studies describe microbial taxonomic shifts associated with geochemical extremes at the ORFRC (Green *et al.*, 2012; Smith *et al.*, 2015; Hemme *et al.*, 2015; Li *et al.*, 2018). For example, *Rhodanobacter* often dominate the microbial community in the most contaminated wells with groundwater pH below 4, nitrate concentrations above 5 mM, and uranium concentrations above 2.5 micromolar (Green *et al.*, 2012; Hemme *et al.*, 2016; Smith *et al.*, 2015; Li *et al.*, 2018). *Rhodanobacter* isolates from the ORFRC, and other environments, can tolerate low pH and possess genes predicted to be involved in metal tolerance (Prakash *et al.*, 2012; Green *et al.*, 2012; van den Heuvel *et al.*, 2010). Thus, although neutral effects can influence microbial communities, the selective pressure of inorganic ion toxicity is likely a dominant factor that influences microbial community composition at this site. However, it is unknown precisely which geochemical parameters favor the growth of *Rhodanobacter* at the ORFRC.

In this study, using a combination of laboratory and field data, we tested the hypothesis that *Rhodanobacter* at the ORFRC are selectively enriched in the most contaminated wells because they are resistant to a specific subset of toxic inorganic ions. Leveraging geochemical and microbial community data from a field survey, we identified 15 inorganic ions that are correlated with elevated relative abundance of *Rhodanobacter*. Next, we used high-throughput cultivation to identify 8 of these ions that (a) are selectively toxic to sensitive microbial isolates from uncontaminated wells at the ORFRC versus resistant *Rhodanobacter* from contaminated wells, and (b) reach concentrations likely to limit the growth of the sensitive isolates, but not *Rhodanobacter*. To validate our results we confirmed the toxicity of mixtures of the selective ions at concentrations representative of the most contaminated wells against ORFRC bacterial isolate collections. Our results clarify which toxic ions are dominant selective pressures on microbial community composition at this contaminated field site, and not merely correlated with taxonomic shifts. Our general approach can be similarly applied to other environments and microbial communities to identify causal factors that influence the relative fitness of microbial sub-populations.

## Materials and Methods

### Comparison of relative abundance data for *Pseudomonas* and *Rhodanobacter* with field concentrations of inorganic ions

Field concentrations of inorganic ions and 16S rDNA sequencing data for different wells at the ORFRC were obtained from the supplementary dataset and through personal communication with the authors of the previous field survey publication (Smith *et al.*, 2015). Spearman correlations between field parameters and microbial taxa were calculated in R. We tested whether each pair of ions (excluding self pairs) was significantly correlated, and P values were converted to false-discovery rates with the p.adjust method. Separately, we tested whether each ion was correlated with any of the top 8 genera and converted those P values to false-discovery rates.

### Inorganic ion arrays

Aqueous solutions of 80 inorganic ions in 96 well format were prepared to capture a wide range of elements from the periodic table in a variety of redox states and to test the influence of chelation on toxicity. Sodium and chloride salts were chosen as counter-ions whenever possible so that sodium chloride could serve as a control for salt stress. All chemicals were purchased from Sigma-Aldrich (St Louis, Mo, USA).

While the oxidation state of most ions was not measured in the field survey, multiple oxidation states are represented in our inorganic ion arrays. Our ion arrays contain multiple oxidation states for an element when those oxidation states are soluble and stable in aerobic, neutral pH aqueous stock solutions (Table S2). Insoluble compounds are less likely to impact microorganisms. Thus, unless we report multiple oxidation states, we posit that the inorganic ion salt in our array is in the dominant soluble oxidation state in ORFRC groundwater samples. Thus, in figures, we represent U(VI) as U, Mn(II) as Mn, etc.

The arrayed compounds were prepared at concentrations close to the solubility limit or at concentrations that we expected would likely be inhibitory to N2E2 (Carlson *et al.*, 2015; Price *et al.*, 2018). Nitrilotriacetic acid (NTA) complexes were prepared by mixing equimolar inorganic ion and NTA. Sterile stock solutions were prepared using 50 mL Steriflip filter units (EMD Millipore, Hayward, CA, USA), transferred to 15 mL conical tubes (BD Biosciences, San Jose, CA, USA), and then a Freedom Evo liquid handling robot (Tecan Group Ltd., Männendorf, Switzerland) was used to transfer 1 mL of stock solution into deep-well 96 well plates (Costar, Thermo Fisher Scientific, Waltham, MA, USA) in the arrayed layout reported in Supplementary Dataset, Table S1.

For growth assays, a Biomek FxP (Beckman Coulter, Indianapolis, IN, USA) liquid handling robot was used to serially dilute stock solutions across the 4 quadrants of three 384-well flat bottom transparent microplates (Costar, Thermo Fisher Scientific) and to add control compounds in a checkerboard pattern in columns A, B, G and H. Positive controls were chloramphenicol (0.2 g/L). Negative controls were water. Each well in the 384 well assay plates contained a final volume of 40 μL aqueous solution. Stock and assay plates were sealed with foil seals (Thermo Fisher Scientific) and stored at −80 °C and thawed 48 hours prior to inoculation. For anaerobic cultures, assay plates were kept unsealed in an anaerobic chamber (COY, Grass Lake, MI, USA) for 48 hours prior to inoculation.

### Media and cultivation conditions

*Pseudomonas fluorescens* FW300-N2E2 was isolated from groundwater from background wells up-gradient from the contaminated area at the ORNL FRC (Thorgersen *et al.*, 2015). *Rhodanobacter sp.* FW104-10B01 (10B01) was isolated from a well within the contaminant plume by direct plating on R2A (R2A, HiMedia, Mumbai, India) agar plates and aerobic growth at 30 °C. *Pseudomonas fluorescens* FW300-N2E2 was recovered from −80 °C freezer stocks in LB aerobically. *Rhodanobacter sp.* FW104-10B01 was recovered from −80 °C freezer stocks in R2A media aerobically. Overnight cultures were washed three times with chemically defined media lacking vitamins, minerals and carbon sources before resuspension in media for dose-response assays. Chemically defined media and KB media recipes are provided in the Supplementary Dataset, Table S2. For anaerobic, nitrate-reducing growth assays, 10 mM nitrate was added to LB media and stored in an anaerobic chamber for 1 week prior to inoculation. N2E2 does not grow fermentatively under anaerobic conditions in LB. For pH profiles, R2A was amended with 30mM HOMOPIPES (pH 3, pH 4), MES (pH 5, pH 6), or PIPES (pH 7) (Buffer salts from Sigma-Aldrich). For all dose-response screens, microbial cells were added to assay plates in an anaerobic chamber (Coy) with a 10% CO_2_:5% H_2_:85% N_2_ atmosphere using a Liquidator 96 (Mettler-Toledo, Oakland, CA, USA). 40 μL of microbial cultures at an optical density (OD 600) of 0.04 in 2x media was added to 384 well flat bottom transparent assay microplates (Costar) containing 40 μL of aqueous compound stocks to obtain a final OD 600 of 0.02, and a final volume of 80 μL. Aerobic assay microplates were sealed with BreathEasy seals (E&K Scientific, Santa Clara, CA, USA) and grown in a Multitron (Infors, Bottmingen, Switzerland) shaker/incubator at 700 rpm at 30 °C. Anaerobic assay microplates were sealed with transparent plate seals (Thermo Fisher Scientific) and grown in the anaerobic chamber without shaking. Growth was monitored by measuring OD 600 with a Tecan M1000 Pro microplate reader (Tecan Group Ltd., Männendorf, Switzerland) using the iControl software package and exported to a Microsoft Excel spreadsheet. A 24-hour timepoint was chosen for most dose-response analysis as this timepoint represents early stationary phase for *Pseudomonas fluorescens* FW300-N2E2 for the media conditions tested. *Rhodanobacter sp.* FW104-10B01 grows more slowly in R2A than does N2E2 and therefore we calculated IC_50_s based on growth at 48 hours for comparisons between N2E2 and 10B01. All growth assays were carried out in duplicate or triplicate and were repeated on different days with similar results.

### Dose-response analysis

Microplate absorbance reads were uploaded to an in-house growth curve database using custom scripts. Data analysis for dose-response inhibition experiments was carried out using the drc R package (Ritz and Streibig, 2005; Ritz *et al.*, 2015) and curves were fit to a standard inhibition dose-response curve to generate an IC_50_ value. In some cases, compound precipitation in growth media impacted absorbance reads. To account for these cases, we removed microplate wells with initial absorbance reads greater than 1.2-fold above control wells from the IC_50_ calculation. Positive and negative controls were used to determine % inhibition values relative to controls and all fitting was carried out on relative inhibition dose-response curves. 95% confidence intervals are reported and all IC_50_s are calculated from at least two biological replicate dose-response curves. For compounds that did not inhibit, or always inhibited greater than 50% across the range of concentrations we screened, we indicate the highest concentration or lowest concentration screened in the Supplementary Dataset, Table S5. Comparisons between growth conditions to assess compounds with differential inhibitory potency are for cultures inoculated on the same day. We define compounds to be selective if the ratio between the IC_50_s (selectivity index, SI) is greater than 2-fold and the 95% confidence intervals do not overlap. For uranium and pH, dose-response data was analyzed using GraphPad Prism 7 (GraphPad Software Inc., La Jolla, CA, USA) and ANOVA was used to compare fits between growth conditions to confirm that the differences between fits for selective compounds were statistically significant. In support of this approach, for replicate dose-response experiments with rich media aerobic cultures of N2E2 (Luria-Bertani broth, LB/aerobic), all compounds display similar inhibitory potencies with overlapping 95% confidence intervals (Figure S1). IC_50_s with confidence intervals are reported in the Supplementary Dataset, Table S5.

### Arrayed isolate collection

Isolates were obtained from groundwater samples from the ORFRC using a variety of aerobic and anaerobic enrichment strategies (Table S6) but all isolates are capable of growth in R2A or LB media aerobically. The arrayed culture collection was prepared from single colony picks and grown in LB or R2A overnight before cryo-preservation in 25% glycerol. Strains were identified using 16S Sanger sequencing and are indicated in Supplemental Dataset, Table S4. To identify abundant sequence types in 16S data from the site (Smith et al 2015), we downloaded the reads from MG-RAST (https://www.mg-rast.org/linkin.cgi?project=mgp8190). We used usearch (https://www.drive5.com/usearch/) to filter out reads with more than one expected error; we extracted the region between the primers (the V4 region) with a custom perl script; and we used usearch to count the abundance of each sequence type in each sample. Arrayed isolates were recovered in R2A in 96 deep well blocks (Costar) aerobically with shaking at 700 rpm in an Infors Multitron plate shaker/incubator. Overnight cultures were centrifuged to pellet cells. Cells were resuspended in buffered R2A media in the presence of inorganic ions at the average concentrations observed in wells with >5% *Rhodanobacter* at an OD600 between 0.01 and 0.06 for sensitivity assays. As mentioned above, 30 mM pH buffers were used for pH profiles. IC_50_s were not affected by addition of 30 mM PIPES to R2A. An isolate was scored as inhibited in a given condition if growth was at least 2 standard deviations below the average of 4 control no stress cultures.

## Results

### Identification of inorganic ion parameters correlated with high relative abundance of *Rhodanobacter* at the ORFRC

We analyzed data from a field survey in which the concentrations of 30 inorganic ion parameters, including pH, were measured in groundwater from 93 wells at the ORFRC (Smith *et al.*, 2015) (Supplemental Table S3). Many of these ions are known to be elevated in the contaminated wells and are associated with microbial community taxonomic shifts (Smith *et al.*, 2015; Li *et al.*, 2018). In this dataset, we found that low pH is significantly positively correlated (false discovery rate <5%) with high concentrations of 19 inorganic ion parameters including U, Al, Mn, Ni, Co, Zn, Cd, Ca, NO_3_^−^, K, Pb, As, Cr, Mg, Be, Ga, Fe, SO_4_^2−^ and Cl (Figure 1A, Supplemental Table S4).

**Figure 1.**
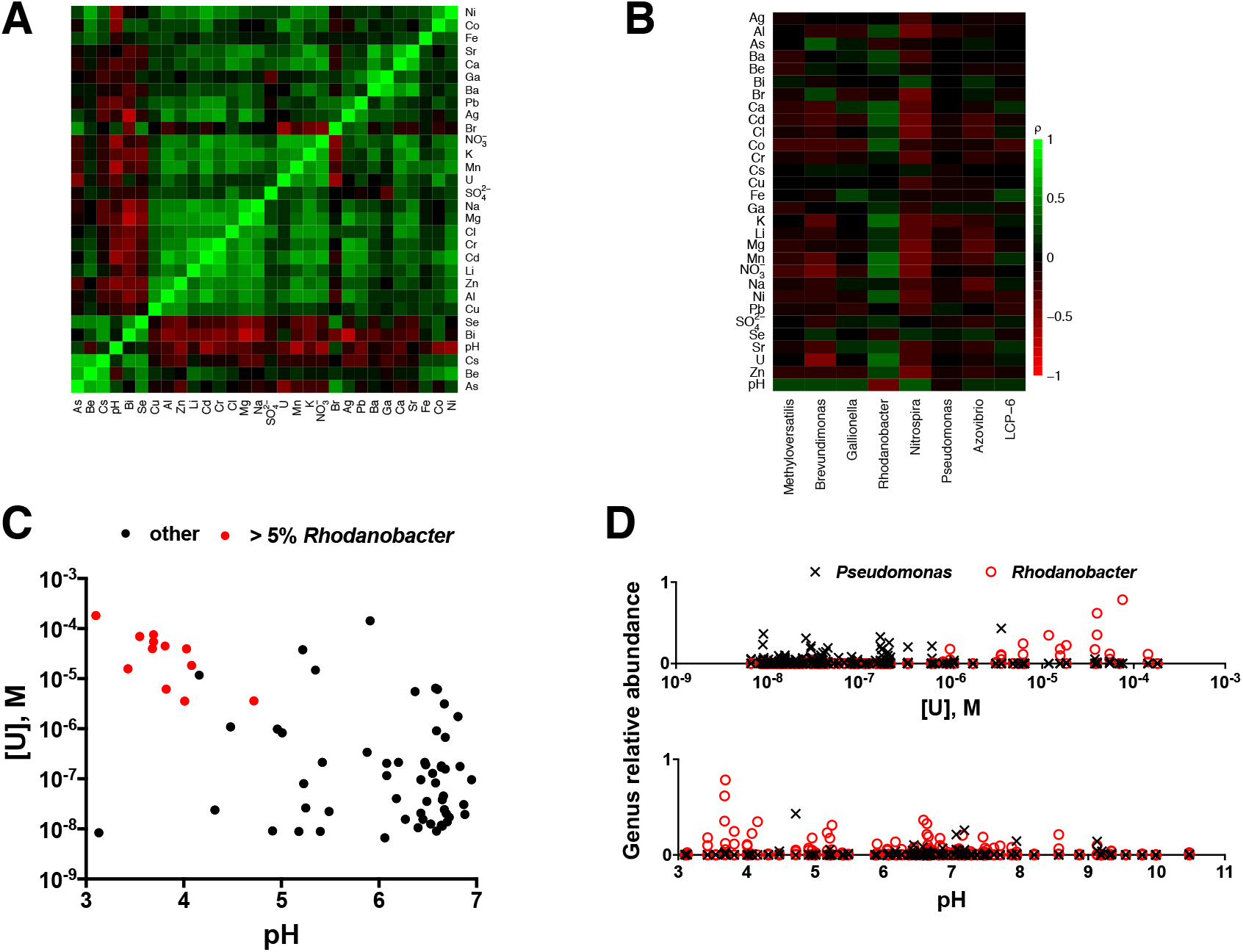
Field correlations between inorganic ions and dominant bacterial taxa at the ORFRC. **A**. Heatmap of Spearman’s rank correlations (ρ) between inorganic ion concentrations in groundwater samples from a field survey at the ORFRC (Smith *et al.*, 2015). Legend is the same as in B. **B**. Heatmap of Spearman’s rank correlations between ion concentrations and the relative abundances of the 8 most abundant bacterial genera. **C**. Concentrations of U and pH in groundwater samples from wells at the ORFRC. Symbols are colored red if the sample also contains >5% *Rhodanobacter*. **D**. Relative abundances of *Rhodanobacter* (red circles) and *Pseudomonas* (black x’s) plotted versus concentrations of U and pH in groundwater samples from the ORFRC.

We also analyzed 16S rDNA amplicon sequences that were collected alongside the geochemical data. We found that the relative abundance of *Rhodanobacter* is significantly positively correlated (false discovery rate <5%) with low pH and high NO3^−^, U, Mn, Al, Co, Zn, Cd, Ni, Ca, Sr, K, Ba, SO_4_^2−^ and Cl (Figure 1B, Table S4), while the other ions are not strongly correlated with *Rhodanobacter* (Figure 1B, Table S4). We also examined correlations for the other 7 genera that are on average most abundant in the field survey dataset. There are few strong correlations between high abundances of these other genera and high concentrations of any of the ions (Figure 1B, Table S4).

*Rhodanobacter* are occasionally observed in uncontaminated wells, but often dominate the microbial community in the most contaminated wells with low pH and high concentrations of U (Figure 1C) and other inorganic ions (Figure S2). In fact, all of the wells with >5% *Rhodanobacter* are the most contaminated wells with both high U concentrations and low pH (Figure 1C). Thus, *Rhodanobacter* are an indicator taxon for contamination at the ORFRC (Smith *et al.*, 2015; Green *et al.*, 2012). There is evidence that *Rhodanobacter* isolates from the ORFRC are acid and metal tolerant (Hemme *et al.*, 2016; Green *et al.*, 2012; Prakash *et al.*, 2012), and we postulated that some, but not all, of the inorganic ions that are correlated with high relative abundance of *Rhodanobacter* wells are selective pressures that limit the growth of less resistant microbial taxa, in the most contaminated wells.

### Identification of inorganic ions selectively inhibitory to a sensitive *Pseudomonas* isolate versus a resistant *Rhodanobacter* isolate

We sought to identify which inorganic ions are selectively permissive for ion-resistant *Rhodanobacter* and restrict ion-sensitive taxa in the most contaminated wells. *Pseudomonas*, though abundant in the field survey dataset (Table S3), are rarely found at high abundance in the presence of high concentrations of inorganic ions and low pH (Figure 1B, 1D; Figure S2). As an exemplar ion-sensitive isolate, we selected a *Pseudomonas* from an uncontaminated well (up-gradient from the contaminant plume), *Pseudomonas fluorescens* FW300-N2E2 (N2E2). We chose N2E2 because previous work indicated that this strain is relatively sensitive to some transition metals (Price *et al.*, 2018), and, as such, is likely representative of many ion-sensitive isolates from uncontaminated wells at the ORFRC.

Because low pH and uranium are the major contaminants at the site, we measured the pH profile and quantified the inhibitory potency of uranium against a *Rhodanobacter* isolate from one of the most contaminated wells, *Rhodanobacter* sp. FW104-10B01 (10B01) and against N2E2. For the purposes of the N2E2/10B01 comparisons, both strains were grown in R2A media to ensure that ion activity would be identical in the growth media used for the comparisons. Consistent with these ions being selectivity determinants, we found that 10B01 is more resistant than N2E2 to low pH (Figure 2A) and to U(VI) (Figure 2B).

**Figure 2.**
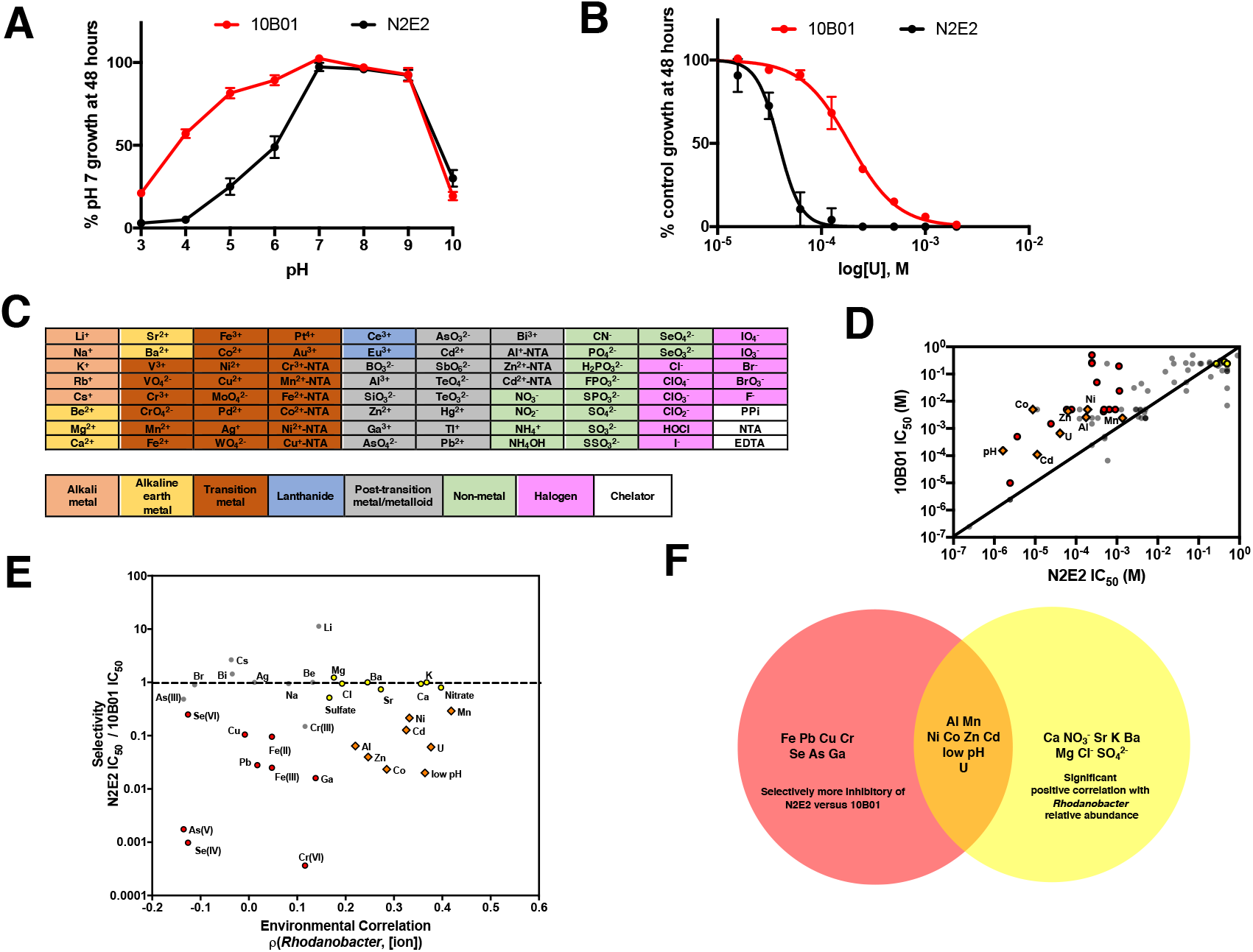
Comparisons of inorganic ion IC_50_s between a sensitive *Pseudomonas* isolate and a resistant *Rhodanobacter* isolate. **A**. pH profiles for *Pseudomonas fluorescens* FW300-N2E2 (N2E2) and *Rhodanobacter sp.* FW104-10B01 (10B01). **B**. Dose-response curves for inhibition of N2E2 and 10B01 by U (uranyl acetate, U(VI)). **C**. The arrayed collection of 80 inorganic ion ions used in dose-response assays to measure inorganic ion toxicity. For some cations, nitrilotriacetic acid (NTA) complexes were prepared. Other abbreviations are PPi (pyrophosphate) and EDTA (ethylenediamine tetraacetic acid). **D**. Comparisons of the IC_50_s for N2E2 and 10B01. Symbols are orange diamonds and labeled if the inorganic ion is both selectively inhibitory of N2E2 and significantly positively correlated with increased *Rhodanobacter* abundance at the ORFRC. Symbols are yellow circles if the inorganic ion is significantly positively correlated with *Rhodanobacter* relative abundance but is not selectively inhibitory of N2E2. Symbols are red circles if the inorganic ion is representative of a geochemical parameter measured in the field survey and is selectively inhibitory of N2E2. pH is plotted as [H_3_O+]. Diagonal line represents the line of equivalence between IC_50_s for N2E2 and 10B01. **E**. Comparisons of the N2E2/10B01 IC_50_ ratios (Selectivity) and the Spearman’s rank correlations for *Rhodanobacter* relative abundance and ion concentrations (Environmental Correlation, *ρ(Rhodanobacter*, [ion]). Coloring is as in C. Horizontal line represent equivalence between IC_50_s for N2E2 and 10B01. **F**. Venn diagram representing the compounds that are either selectively inhibitory of N2E2 versus 10B01 (red), significantly positively correlated with increased *Rhodanobacter* abundance in the field survey (yellow) or both (orange).

To identify other inorganic ions to which 10B01 is resistant compared to N2E2, we determined the inhibitory potency of a panel of 80 inorganic ions (Figure 2C) against both isolates grown in R2A media (Figure 2D). We chose these ions because they likely represent the dominant oxidation states of most elements found in groundwater, or when multiple oxidation states are common, we included both. We quantify inhibitory potency as the concentration required to inhibit growth to 50% of uninhibited control cultures (IC_50_), and we quantify selectivity as the ratio between two IC_50_s, (e.g. N2E2 IC_50_/10B01 IC_50_) (Supplemental Dataset, Table S5). We consider a compound selective if the 95% confidence intervals of the dose-response curves for the two organisms do not overlap and the IC_50_s differ by more than a factor of 2. By these criteria, alongside low pH and U(VI), 10B01 is more resistant than N2E2 to Al(III), Mn(II), Ni(II), Co(II), Zn(II), Cd(II), Cr(VI), Fe(II), Fe(III), Pb(II), Se(IV), Se(VI), As(V), Ga(III), and Cu(II) (Figure 2D, Figure 2E). While heavy metal resistance genes are present in *Rhodanobacter* genomes (Hemme *et al.*, 2016; Green *et al.*, 2012; Prakash *et al.*, 2012), similar resistance determinants are present in the genomes of N2E2 and other *Pseudomonas* (Thorgersen *et al.*, 2017; Price *et al.*, 2018). Our results are the first experimental demonstration that these specific ions can select for *Rhodanobacter* similar to 10B01 over less tolerant organisms from the ORFRC such as *P. fluorescens* FW300-N2E2.

Of the ions to which 10B01 is resistant and for which we have measurements of the corresponding element in the field data, low pH and high U(VI), Mn(II), Al(III), Zn(II), Ni(II), Co(II) and Cd(II) are positively correlated with *Rhodanobacter* relative abundance in the field survey dataset (Figure 2E, 2F). All of these are likely the dominant soluble oxidation state measured in the field survey samples. In contrast, N2E2 and 10B01 are similarly resistant to the alkali earth metal cations Sr(II), Ba(II), Mg(II), K(II), and the anions Cl^−^ and SO_4_^2−^ and thus, these ions are less probable selectivity determinants. These ions are far less toxic, with IC_50_s for both isolates several orders of magnitude above those of the transition metals (Supplemental Dataset Table S5). Finally, 10B01 is more resistant to the metal cations Fe(II), Fe(III), Pb(II), Ga(III), and Cu(II) and oxyanions Se(IV), Se(VI), As(V), Cr(VI) relative to N2E2, but none of these ions are positively correlated with increased *Rhodanobacter* abundance, and thus, these ions are probably less important in selecting for *Rhodanobacter* at the ORFRC. These ions could be selectivity determinants in some locations but are not dominant selective pressures (Supplemental Dataset, Table S3).

### Identification of inorganic ion toxicity thresholds in the field that can limit a sensitive *Pseudomonas* isolate and favor a resistant *Rhodanobacter* isolate

A selective ion must reach concentrations in the field that, while permissive for *Rhodanobacter*, limits the survival of other taxa. Thus, we compared the IC_50_s measured against N2E2 with ion concentrations in the most contaminated wells (Figure 3). We include wells in the set of “the most contaminated wells” if they have >5% *Rhodanobacter.* As, noted above, all of the most contaminated wells have low pH and elevated concentrations of U and other inorganic ions (Figure 1C, Figure S3).

**Figure 3.**
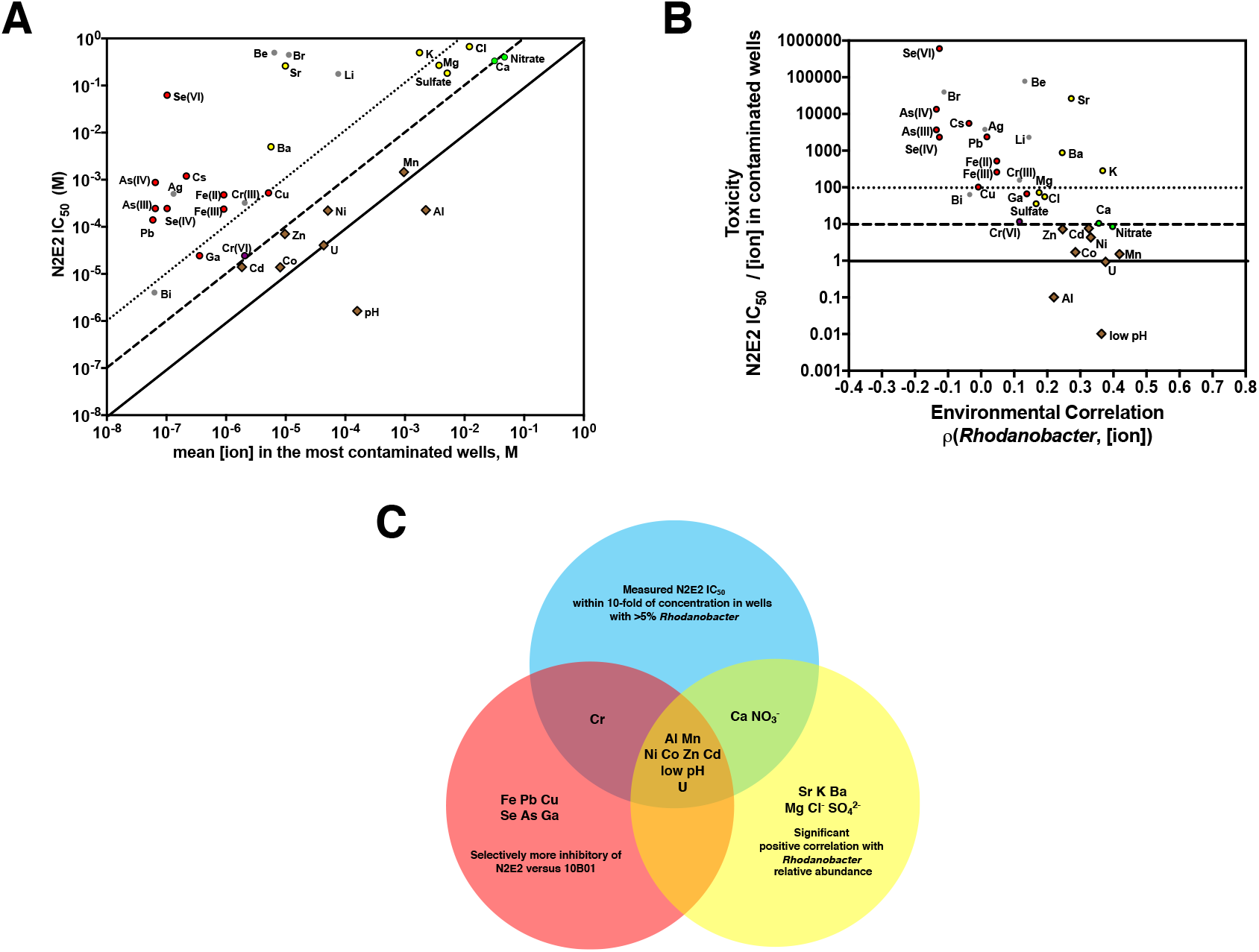
Comparisons of inorganic ion IC_50_s with field concentrations. Symbol coloring throughout as in panel C. **A**. *Pseudomonas fluorescens* FW300-N2E2 (N2E2) IC_50_s compared with mean concentration of inorganic ion parameters in wells with >5% *Rhodanobacter.* pH is plotted as [H_3_O+]. Diagonal lines represent N2E2 IC_50_s equivalent to (solid), 10-fold below (dashed) and 100-fold below (dotted) the mean [ion] in the most contaminated wells. **B**. Comparisons of the ratios of N2E2 IC_50_/average field ion concentration in the most contaminated wells and the Spearman’s rank correlations between *Rhodanobacter* relative abundance and ion concentrations. **C**. Venn diagram of (i) inorganic ion parameters selectively inhibitory of N2E2 versus 10B01, (ii) significantly positively correlated with increase *Rhodanobacter* abundance in the field survey, (iii) within 10-fold of the average field concentration in wells with >5% *Rhodanobacter*. Horizontal lines are marked as in 3B.

The IC_50_s against N2E2 for low pH, U(VI), Mn(II), Al(III), NO3^−^, Ca(II), Cd(II), Co(II), and Zn(II) are all within 10-fold of the concentrations in contaminated groundwater wells with >5% *Rhodanobacter* (Figure 3A). Strikingly, all of these ions are significantly positively correlated with *Rhodanobacter* relative abundance in the field (Figure 3B), and, with the exception of Ca(II) and NO_3_^−^, are selectively inhibitory of N2E2 versus 10B01 (Figure 2). In general, there is good agreement between how strongly an ion is correlated with *Rhodanobacter* abundance and how close the N2E2 IC_50_ is to the field concentration (Figure 3B). For example, while Pb(II) and Al(III) have similar N2E2 IC_50_s, the Al concentration is almost 5-orders of magnitude higher, and only Al is positively correlated with *Rhodanobacter* relative abundance. Thus, Al, but not Pb, is likely to impact N2E2 range at the ORFRC (Figure 3A). On the other hand, Sr, K, Ba, Mg, Cl^−^ and SO_4_^2−^ are positively correlated with increased *Rhodanobacter* relative abundance, but are not selectively inhibitory of N2E2 versus 10B01 (Figure 2, Figure 3C), and their concentrations in the contaminated wells are more than 10-fold below the N2E2 IC_50_ (Figure 3B).

The oxidation state of most ions was not measured in the field survey, but for Fe, Te, Se, Cr and As the inhibitory potency varies based on oxidation state (Figure 2E, 3A). For example, while the Cr(VI) IC_50_ is close to the field concentration of Cr, Cr(III) is much less toxic (Figure 3C). Thus, although Cr is not strongly correlated with *Rhodanobacter*, it may be a selective pressure on organisms like N2E2 in the contaminated wells if the dominant oxidation state is Cr(VI).

### Quantifying the impact of growth medium changes on inorganic ion toxicity

Based on our initial dose-response assays with N2E2 and 10B01, we identified low pH and high U(VI), Mn(II), Al(III), Cd(II), Zn(II), Co(II), and Ni(II) as (a) correlated with *Rhodanobacter* relative abundance in the field (Figure 1), (b) selectively inhibitory of N2E2 versus 10B01 (Figure 2) and (c) reaching average concentrations within a factor of 10 of the N2E2 IC_50_ in the ORFRC wells with >5% *Rhodanobacter* (Figure 3). However, the growth conditions in the aquifer vary widely from the aerobic growth conditions in R2A medium in which we measured ion inhibition of 10B01 and N2E2. To measure the influence of this variation on ion toxicity, we measured N2E2 IC_50_s under different growth conditions (Figure 4).

**Figure 4.**
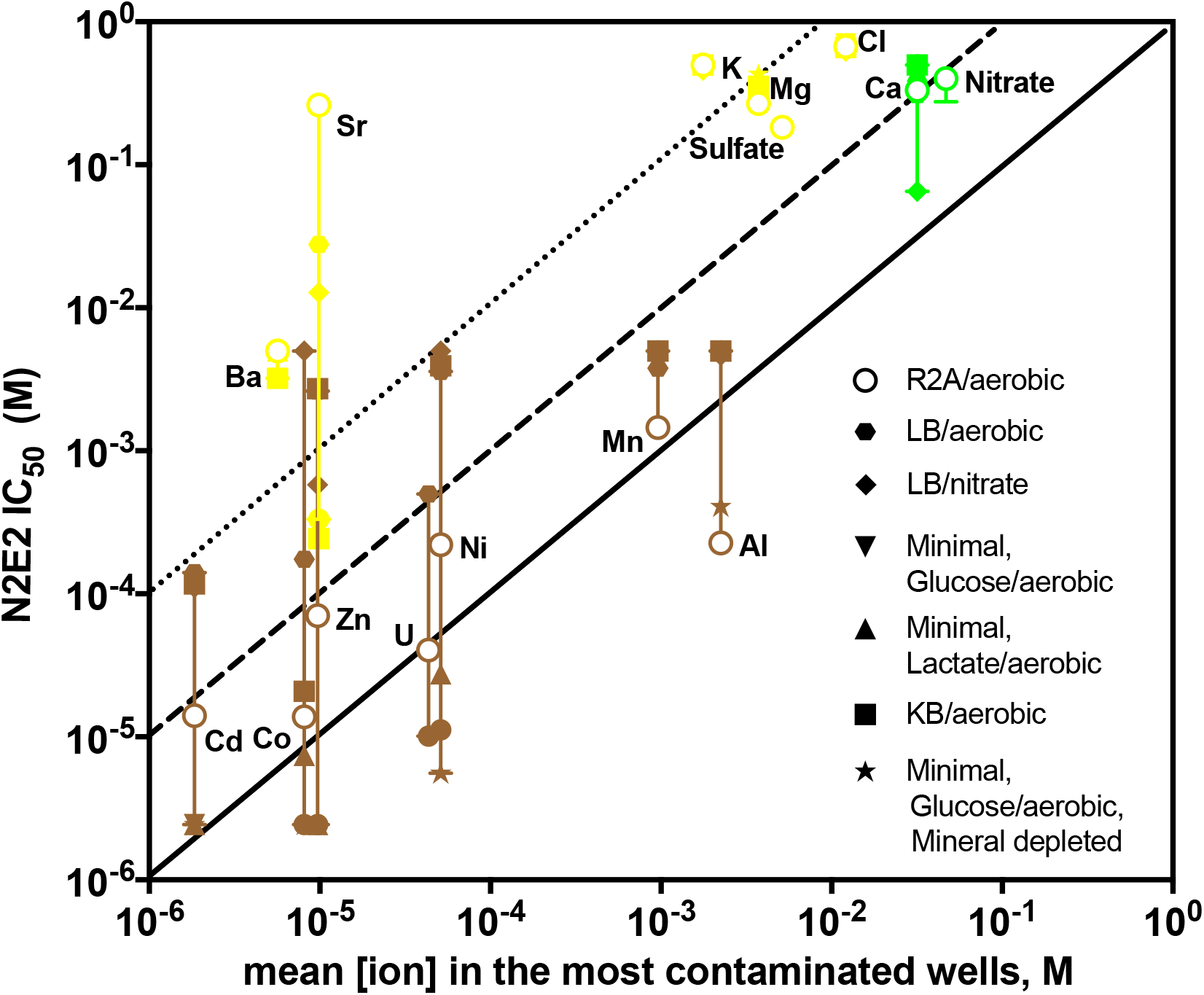
Impact of growth media conditions on inorganic ion inhibitory potency. *Pseudomonas fluorescens* FW300-N2E2 (N2E2) IC_50_s measured for inorganic ion parameters correlated with high *Rhodanobacter* abundance. IC_50_s in each growth condition are indicated by symbols in the legend, and error bars represent the highest and lowest IC_50_s measured in any growth medium and compared with average concentrations in the most contaminated wells. Coloring is as in Figure 3C. Diagonal lines are marked as in 3B.

Because metal toxicity is controlled by free metal activity (Nowack and VanBriesen, 2005; Leštan *et al.*, 2008; Hughes and Poole, 1991) we measured the impact of varying organic carbon content on ion toxicity. The metal-binding capacity of groundwater samples from the ORFRC is low, with dissolved organic carbon ranging between 10 and 100 mg/L (McCarthy *et al.*, 1993; Drewes and Fox, 1999; Davis, 1984; Smith *et al.*, 2015; Wu *et al.*, 2018). On the other hand, organic carbon in forest soils, such as those at the ORFRC, can be up to 2% by weight (Osman, 2013). Thus, depending on the depth in the aquifer, the carbon content may dramatically alter inorganic ion IC_50_s. The microbial community in well water may represent microorganisms that grew at any location in this carbon gradient and were carried by groundwater flow to wells. Thus, many ORFRC microorganisms may experience dramatic variation in organic carbon concentrations. Furthermore, organic carbon content at the ORFRC has been altered artificially through the injection of carbon sources such as emulsified vegetable oil or ethanol (Li *et al.*, 2018; Watson *et al.*, 2013). In these cases, local carbon concentrations may exceed that found in soils or in rich microbial growth media.

Aside from the influence of organic carbon on inorganic ion toxicity, different metabolic states of bacterial cells are differentially susceptible to antimetabolic ions. For example, the enzymes of different carbon utilization pathways are differentially susceptible to inhibitory ions (Ong *et al.*, 2015; Klemperer, 1950; Benabe *et al.*, 1987). Under aerobic conditions, toxic reactive oxygen species are generated by redox active metals (Lemire *et al.*, 2013; Strlič *et al.*, 2003), while under nitrate-reducing conditions, various oxyanion substrates of nitrate reductases are differentially toxic depending on whether they are detoxified or converted to a more toxic form by the enzyme (Oremland and Stolz, 2000; Oremland and Capone, 1988; Garbisu *et al.*, 1996). Under trace metal depleted conditions, toxic ions can compete with nutritional metals for uptake systems (Lemire *et al.*, 2013; Braud *et al.*, 2009; Wichard *et al.*, 2009; Teitzel and Parsek, 2003; Enoch and Lester, 1972).

To systematically investigate condition-dependent ion sensitivity, we measured the impact on the toxicity of inorganic ions of varying the carbon source, terminal electron acceptor, trace metal availability and iron availability (Figure 4B). We observed many intriguing changes in IC_50_s by varying growth conditions, and likely mechanisms are discussed in Supplemental Note 1. We focused on identifying conditions in which U(VI), Mn(II), Al(III), Cd(II), Co(II), Zn(II) and Ni(II) are less toxic to N2E2 or Sr(II), Ba(II), Mg(II), Cl^−^, SO_4_^2−^ and K(II) are more toxic, as this might alter our list of selectivity determinants that favor *Rhodanobacter* at the field site. Thus, for this subset of ions, we compared the maximum and minimum N2E2 IC_50_s from all growth conditions against field concentrations (Figure 4).

The inhibitory potency of Mn(II), Al(III), Ca(II) and NO3^−^ against N2E2 are not greatly affected by changes in growth conditions (Figure 4, Figure S4). Only dramatic increases in growth medium organic carbon alleviates the toxicity of U(VI), Cd(II), Co(II), Zn(II) and Ni(II) (Figure 4A, Figure S4, Table S5) such that that these ions will not be inhibitory to N2E2 in the contaminated wells at high carbon concentrations. Conversely, in minimal medium, at low carbon concentration, the N2E2 IC_50_ for Sr decreases to ~30-fold above the average concentration in the wells with >5% *Rhodanobacter* (Figure 4, Table S5). Thus, although we find it unlikely, we cannot eliminate the possibility that there are carbon-limited environments at the ORFRC where Sr(II) toxicity impacts microbial community composition. IC_50_s for Ba(II), Mg(II), Cl^−^, SO_4_^2−^ and K(II) are not greatly affected by changes in growth media composition (Figure 4), and thus, it is unlikely that these ions are directly toxic to ion-sensitive organisms such as N2E2 at any location at the ORFRC under any conditions (Figure 4).

While 10B01 does not grow in our chemically defined medium, and we could not evaluate as many growth conditions as with N2E2, comparison of the *Rhodanobacter* R2A IC_50_s with field ion concentrations is informative (Figure S3). For Al(III), Mn(II), Ca(II), NO3^−^ and low pH, the 10B01 IC_50_s are within a factor of 10 of the concentrations in the most contaminated wells. Thus, the free ion activity in the contaminated wells permissive for *Rhodanobacter* is not likely to be much higher than in R2A.

Additionally, *Rhodanobacter* are often implicated in denitrification in the most contaminated wells (Li *et al.*, 2018; Hemme *et al.*, 2010; 2016; 2015), and we were able to compare 10B01 IC_50_s between aerobic and nitrate reducing conditions (Table S5). As with N2E2, we observed some differences in ion toxicity to 10B01 depending on the terminal electron acceptor (Table S5), but not for U(VI), Mn(II), Al(III), Co(II), Cd(II), Ni(II) and Zn(II). These results are consistent with the view that this set of selective ions are important regardless of whether nitrate reduction or aerobic respiration is the dominant terminal electron accepting process.

### Confirmation of selectivity determinants for panels of ORFRC isolates

To test if low pH and high U(VI), Mn(II), Al(III), Cd(II), Co(II), Zn(II) and Ni(II) are selective pressures that favor resistant microbial sub-populations in the contaminated wells, we measured the growth of 194 isolates from the ORFRC in the presence of these ions at their average concentrations in the most contaminated wells (Figure 5, Table S3). For these assays, we grew the isolates in R2A medium aerobically and quantified inhibition relative to control cultures grown in the absence of inhibitory ions (Materials and Methods). Of the arrayed isolates, only 8 *Rhodanobacter* from contaminated wells are capable of robust growth in all conditions we tested (Figure 5A). None of the other 186 isolates, including 123 *Pseudomonas* and 63 other bacteria from uncontaminated wells, are resistant to a mixture of U(VI), Mn(II), Al(III), Cd(II), Co(II), Zn(II) and Ni(II) at pH 4 (Figure 5A, Table S6). In contrast, most isolates are resistant to the ions that met fewer of the criteria for a selective ion including Ca (73%), NO_3_^−^ (83%) Cr(III) (75%) and a mixture of all other 19 ions measured in the field survey (72%) (Figure 3A, Figure 5A). These results support our hypothesis that low pH, and high U(VI), Mn(II), Al(III), Cd(II), Co(II), Zn(II) and Ni(II) are dominant selective pressures in the contaminated wells at the ORFRC, while the other 22 ions we considered are less important.

**Figure 5.**
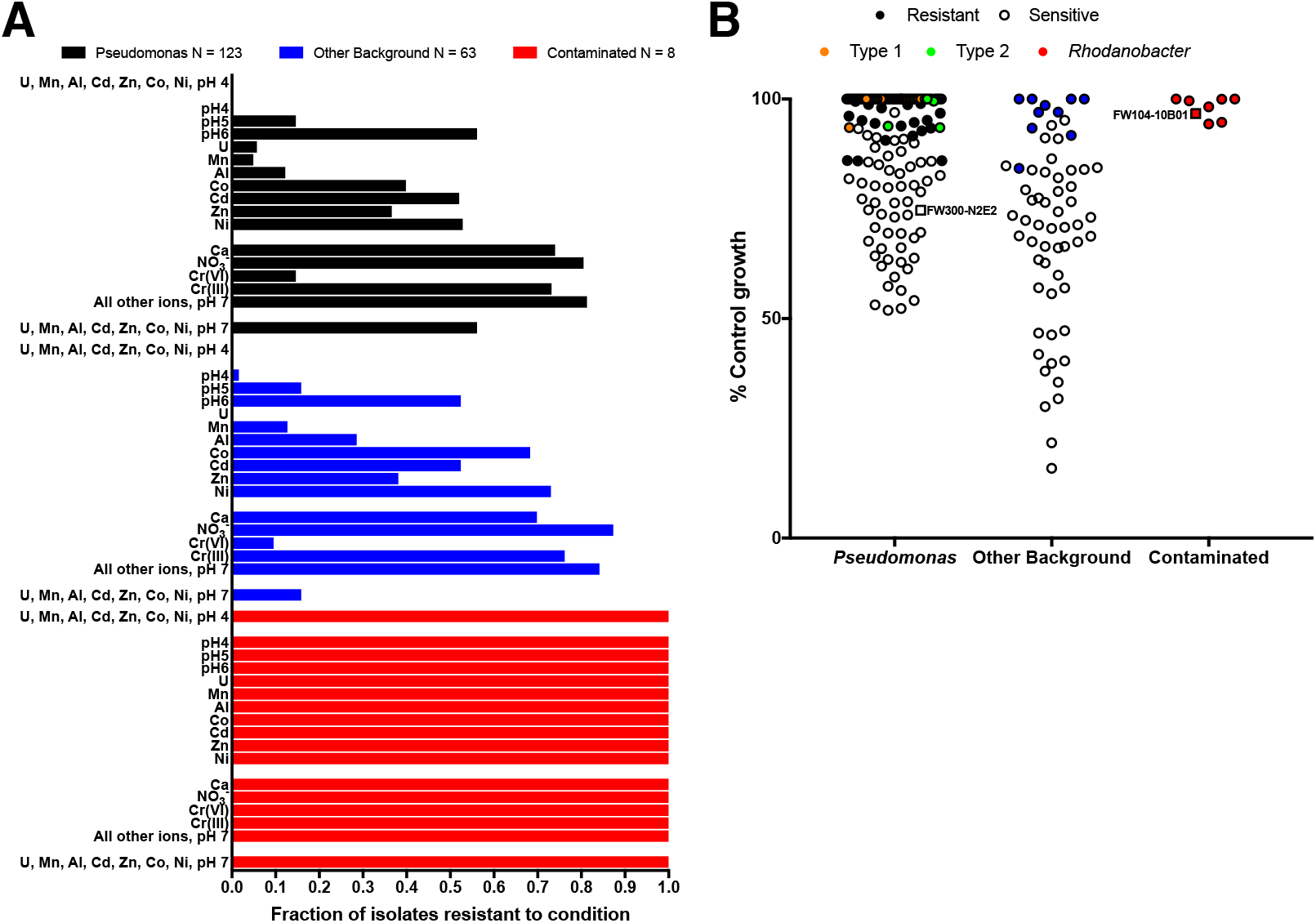
Confirmation of inorganic ion selectivity determinants that favor resistant taxa such as *Rhodanobacter* at the ORFRC. **A**. Fraction of arrayed ORFRC isolates resistant to inorganic ions and their mixtures at the average concentrations in wells with >5% *Rhodanobacter*. Isolates are split into 3 categories (i) *Pseudomonas*, (ii) other background isolates from wells up-gradient from the contaminant plume, and (iii) *Rhodanobacter* isolates from contaminated wells. “All other ions” are the other 19 parameters we considered aside from low pH, U, Mn, Al, Cd, Co, Zn Ni, Ca, NO_3_^−^ and Cr. Unless otherwise indicated, the oxidation state is the same as that in our 80 inorganic ion array (Materials and Methods, Table S1). **B**. Inhibition of *Pseudomonas* isolates, other isolates from background wells and *Rhodanobacter* isolates from contaminated wells by a mixture of U, Mn, Al, Co, Cd, Zn, Ni at pH 7 at the average concentrations in the most contaminated wells. Exact 16S V4 sequence matches with two dominant *Pseudomonas* sub-populations in the contaminated well, PTMW02, are colored orange (Type 1) and green (Type 2).

Some of the ions that are not dominant selective pressures inhibit the growth of some of the more sensitive isolates. Although Ca(II) and NO3^−^ are not selectively inhibitory of N2E2 versus 10B01, they are correlated with high *Rhodanobacter* and the N2E2 IC_50_s are only about 10-fold above the concentrations in the contaminated wells. Thus, it is not surprising that Ca(II) inhibits 27% and NO3^−^ inhibits 17% of the 194 isolates we tested. While Cr is not correlated with high *Rhodanobacter*, Cr(VI) is selectively inhibitory of N2E2 and field concentrations are close to the N2E2 IC_50_ and inhibits 87% of isolates, but Cr(III) is much less toxic and only inhibits 25% (Figure 3).

U, Mn, Al, Cd, Co, Zn and Ni are positively correlated with low pH (Figure 1A, Supplemental Dataset, Table S4), but the concentrations of these metals also vary independently. Thus, we measured the fraction of isolates resistant to each selective ion alone (Figure 5B). Few isolates are resistant to low pH (pH 4, 15%), U(VI) (4%), Mn(II) (8%) and Al(III) (18%), but more isolates are resistant to Cd(II) (51%), Zn(II) (37%), Co(II) (49%), and Ni(II) (60%). Transition metals could be inhibitory to more isolates in conditions with lower organic carbon (Figure 4), but these results demonstrate that the selective ions as isolated parameters can impact the microbial community in the contaminated wells (Figure 5A).

Our results also enable us to classify ORFRC field isolates based on resistance to the selective inorganic ions. We measured resistance to a neutral pH mixture of U(VI), Mn(II), Al(III), Cd(II), Co(II), Zn(II) and Ni(II) and found that all *Pseudomonas* isolates with 16S V4 regions identical to the two dominant sub-populations (Type 1 and Type 2, Figure 5B) in a contaminated well, PTMW02 (Table S3, Table S6), were resistant to concentrations of these metals in the wells with >5% *Rhodanobacter.* These results are consistent with the success of resistant *Pseudomonas* strains in one of the most contaminated wells, especially when pH is closer to neutral. PTMW02, with a pH of 4.78, is has the highest pH of any of the most contaminated wells (Table S3).

### Non-additive interactions between inorganic ions can influence toxicity thresholds

By mixing the selective inorganic ions in formulations representative of the contaminated wells, we gained insights into how complex mixtures of ions interact with microbial populations. For example, we noticed that more isolates (42%) are resistant to the neutral pH mixture of U(VI), Mn(II), Al(III), Cd(II), Co(II), Zn(II) and Ni(II) than to Mn(II) (8%) or U(VI) (4%) alone (Figure 5B). This suggests antagonism, and when we measured the N2E2 IC_50_s for U(VI) in combination with Mn(II), we found the interaction to be antagonistic (Figure S6). Antagonism is known for interactions between U(VI) and metals for inhibition of the aquatic plant, *Lemna aequinoctialis* (Charles *et al.*, 2006) and for *Hydra viridissima* (Hyne *et al.*, 1992), but to our knowledge, this is the first report of such antagonism in bacteria. Although this antagonism has a small impact on microbial fitness compared to other changes in growth conditions, it is important to consider in the context of the complex multi-metal gradients at the ORFRC.

## Discussion

In diverse environments, including the ORFRC, the increased prevalence of ion resistant taxa is positively correlated with increased ion concentrations (Smith *et al.*, 2015). However, in complex ion gradients, the dominant selective ions are difficult to identify. In this study we developed a systematic approach for identifying selectively toxic inorganic ions. We found that low pH and high U(VI), Mn(II), Al(III), Cd(II), Co(II), Zn(II) and Ni(II) are likely selective pressures in the most contaminated wells at the ORFRC because these ions are (a) positively correlated with *Rhodanobacter* relative abundance across the field site, (b) selectively inhibitory of non*-Rhodanobacter* isolates from uncontaminated wells and (c) reach toxic concentrations in the contaminated wells that are inhibitory to most isolates from uncontaminated wells.

Our results point to specific areas where future field measurements will enhance our understanding of the ORFRC. For example, because the toxicity of some ions, (e.g. Cd(II), Zn(II), Co(II) and Ni(II)) varies as a function of growth conditions (e.g. organic carbon content), it is difficult to quantitatively rank the relative importance of each selective ion without more data on carbon source, electron acceptor and nutrient availability at a given location. Thus, future work to define the influence of natural organic matter from the ORFRC (Giller *et al.*, 1998) on ion toxicity will improve our ability to predict ion impact on microbial communities in the aquifer. In addition, our conceptual model of the ORFRC will also be improved with more data on the prevalence and oxidation state of some ions. For example, more data on the redox speciation of Se, Te, As, Cr, and Fe, which vary dramatically in toxicity depending on oxidation state, will enable us to identify locations in the aquifer where these elements might be selective pressures. For some of the most toxic elements (e.g. Hg, Th, Pd, Ce, Cs, Au), the N2E2 IC_50_ was below the lowest concentration we tested in our dose-response assays, but we have limited data on the prevalence of these ions. Mercury (Hg) has been measured in ORFRC groundwater at concentrations of up to 10 nM (Pereira *et al.*, 2010) which suggests that mercury concentrations could limit N2E2 growth at some locations in the field, and mercury efflux systems are observed in *Rhodanobacter* from the most contaminated wells at the ORFRC (Hornberger *et al.*, 1999).

Many studies on the microbial ecology of metal contaminated aquifers focus on the interpretation of taxonomic shifts in terms of probable physiological traits associated with U(VI) bioreduction and immobilization (Li *et al.*, 2018; Watson *et al.*, 2013; Fields *et al.*, 2005; Williams *et al.*, 2013). Future studies at the ORFRC should consider the impact of the toxic ions we identified on microbial metabolic capabilities implicated in controlling U mobility. For, example while *Rhodanobacter* are highly resistant to conditions in the contaminated wells, they are associated with nitrate reduction and U mobilization (Li *et al.*, 2018; Fields *et al.*, 2005; Williams *et al.*, 2013; Hemme *et al.*, 2010). Our results indicate that the conditions in the contaminated wells are permissive for *Rhodanobacter* independent of respiratory state, but other denitrifiers and the Fe(III) or SO_4_^2−^ reducing microorganisms that mediate U reduction and immobilization (Li *et al.*, 2018; Williams *et al.*, 2013) may differ in sensitivity to the toxic ions in the contaminated wells. High-throughput dose-response assays can also be used to identify specific inhibitors of respiratory metabolisms (Carlson *et al.*, 2015; 2017a; 2017b), and targeted interventions such as the addition of chelators (Leštan *et al.*, 2008), or choosing carbon sources and carbon concentrations to minimize metal toxicity (Vanengelen *et al.*, 2010) may improve the efficacy of U bioremediation efforts.

By clarifying which inorganic ions at the ORFRC are likely to influence microbial communities, our results enable future studies to define the biological mechanisms of resistance. Low pH, high NO3^−^ and high U(VI) are commonly referenced as markers of contamination at the ORFRC and all are correlated with high *Rhodanobacter* abundance (Smith *et al.*, 2015; Li *et al.*, 2018; Brooks, 2001; Williams *et al.*, 2013; Thorgersen *et al.*, 2017). While U(VI) and low pH selectively inhibitory of *Pseudomonas*, NO_3_^−^ is not, but may impact community composition as an important terminal electron acceptor. Few studies mention Mn(II) and Al(III) toxicity as important at the ORFRC or focus on understanding the mechanisms of resistance to these metals, but they are important selective pressures in the contaminated wells. Also, metal efflux systems are common in both *Pseudomonas* (Price *et al.*, 2018; Thorgersen *et al.*, 2017) and *Rhodanobacter* (Hemme *et al.*, 2010; 2016; 2015), but the ion specificity of the resistance mechanisms for U(VI), Mn(II), Al(III), Cd(II), Co(II), Zn(II) and Ni(II) is poorly understood.

Finally, our approach could be applied to other environments with elevated concentrations of inorganic ions such as the rhizosphere (Giller *et al.*, 1998), oil reservoirs (Pereira *et al.*, 2010), wastewater (Stepanauskas *et al.*, 2005) or marine sediments (Hornberger *et al.*, 1999). While the ORFRC is an extreme example, there are indications that inorganic ion toxicity can impact microbial populations in many environments (Duxbury, 1985; Fierer, 2017; Gadd, 2010). More broadly, understanding the biogeochemical controls on microbial activity in the environment is a central challenge of environmental microbiology, and we anticipate that further efforts to array biological and geochemical diversity in a format amenable to high-throughput cultivation will enable more rapid and accurate identification of the controls on microbial community composition and activity.

## Acknowledgements

This work was funded by ENIGMA, a Scientific Focus Area Program supported by the U. S. Department of Energy, Office of Science, Office of Biological and Environmental Research, Genomics: GTL Foundational Science through contract DE-AC02-05CH11231 between Lawrence Berkeley National Laboratory and the U. S. Department of Energy. The authors would like to thank members of the ENIGMA Scientific Focus Area for critical feedback and suggestions.

## Supplementary Materials

### Supplementary Dataset with Tables S1-S6 is provided in a separate.xlsx file

**Table S1**. 96 well-layout 80 inorganic ion array plate layout. Compound name, CAS numbers, and stock concentrations in M and mM are indicated.

**Table S2**. Media recipes for King’s B media (KB) and chemically defined minimal media (minimal media) used for growth assays.

**Table S3**. 16S amplicon sequencing data and concentrations of various geochemical parameters identified as possible controls on N2E2 based on our study. Data from Smith, M. B. *et al.* Natural Bacterial Communities Serve as Quantitative Geochemical Biosensors. *mBio* 6, e00326–15–13 (2015) and personal communication with the authors.

**Table S4**. Spearman correlations between relative abundances of the *Pseudomonas* and *Rhodanobacter* genera from 16S amplicon sequencing data and concentrations of various geochemical parameters identified as possible controls on N2E2 based on our study. Data from Smith, M. B. *et al.* Natural Bacterial Communities Serve as Quantitative Geochemical Biosensors. *mBio* 6, e00326–15–13 (2015) and personal communication with the authors.

**Table S5.** IC_50_ concentrations (M) and selectivity indices (SI) for 80 inorganic ions against *Rhodanobacter sp.* FW104-10B01 and *Pseudomonas fluorescens* FW300-N2E2 in different growth conditions.

**Table S6**. Arrayed ORFRC isolate sensitivity to individual components and mixtures of compounds at the average concentrations in groundwater samples with >5%

*Rhodanobacter.* % inhibition relative to controls is reported, and significant inhibition is indicated for cultures inhibited more than 2 standard deviations below the mean of positive control no stress cultures.

### Supplementary Methods

#### Quantification of synergistic inhibition

Synergistic inhibition between inhibitors was assessed using the equation for Fractional Inhibitory Concentration Index (FICI) based on the IC_50_ for each inhibitor A and B in the absence (IC_50_(A), IC_50_(B)) or presence of the other inhibitor (IC_50_(AB), IC_50_(BA)):

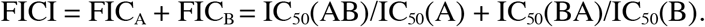

A FICI score < 0.5 implies synergism, whereas a FICI score > 2 implies antagonism. An FICI between 1 and 2 implies indifference, and by convention, the greatest absolute value FICI score is reported (EUCAST, 2000).

European Committee for Antimicrobial Susceptibility Testing of the European Society of Clinical M, Infectious D (2000). EUCAST Definitive Document E.Def 1.2, May 2000: Terminology relating to methods for the determination of susceptibility of bacteria to antimicrobial agents. *Clinical microbiology and infection : the official publication of the European Society of Clinical Microbiology and Infectious Diseases* **6:** 503-508.

### Supplementary Note

#### Insights into variability and mechanisms of inorganic ion resistance from dose-response assays with *P. fluorescens* FW300-N2E2 Organic carbon content

To evaluate how N2E2 IC_50_s are impacted by organic carbon content we measured the inhibitory potency of the 80 inorganic compounds against N2E2 grown in a chemically-defined minimal medium containing ~6g/L organic carbon and in a rich media (LB, Luria-Bertani broth) with ~20 g/L organic carbon (Figure S4B). We observed lower N2E2 IC_50_s in minimal medium relative to rich medium for U and the transition metals Ni, Cd, Zn and Co, such that these parameters are likely not inhibitory to sensitive, background organisms such as N2E2 in carbon rich environments such as organic rich soil horizons (Osman, 2013) or under carbon injection regimes at the ORFRC (Li *et al.*, 2018; Watson *et al.*, 2013), but will be toxic in typical low carbon aquifer environments with measured dissolved organic carbon between 10 and 100 mg/L (McCarthy *et al.*, 1993; Drewes and Fox, 1999; Davis, 1984; Smith *et al.*, 2015). In some cases, rich/minimal IC_50_ ratios are greater than 10^3^, but nitrilotriacetic acid (NTA) complexation alleviated ion toxicity in minimal media (Figure S5, Supplementary Dataset, Table S5). Together, these results are consistent with the view that the ion binding capacity of growth media controls toxicity (Nowack and VanBriesen, 2005; Leštan *et al.*, 2008; Hughes and Poole, 1991).

Similar to the impact of the organic carbon content in growth media, the capacity to form biofilms is likely to give some microorganisms an advantage in the contaminated wells. Biofilm formation by *Pseudomonas* can increase the IC_50_s of inorganic ions by over an order of magnitude, likely due to the ion binding capacity of the extracellular polysaccharide matrix produced by sessile cells (Workentine *et al.*, 2008). However, in no known case does biofilm formation increase the sensitivity of a strain to inorganic ions (Workentine *et al.*, 2008).

#### Organic carbon source

We evaluated how changing the carbon source from glucose to lactate impacts inhibitory potency (Figure S4C). In general, we saw very minor influence of lactate on IC_50_s, aside from a few compounds that may be antimetabolites for the enzymes involved in carbon catabolism. The enzymes of different catabolic pathways have different susceptibilities to inorganic compounds (Ong *et al.*, 2015) . VO_4_^2−^ and Be^2^+ are more inhibitory of glucose grown N2E2 cultures than lactate grown cultures. Both compounds can interfere with phosphate transfer reactions in glycolysis (Benabe *et al.*, 1987; Klemperer, 1950). We did not observe any compounds with greater inhibitory potency under lactate utilizing conditions.

#### Terminal electron acceptor

N2E2, like many Pseudomonads, is capable of complete denitrification of NO3^−^ to N_2_ (Thorgersen *et al.*, 2015b). We determined the IC_50_s of the 80 inorganic compounds on the anaerobic growth of N2E2 with LB as the carbon source and electron donor and 10 mM nitrate as the sole electron acceptor (Figure S4D). In general, the redox state for Te, Se, As and Cr has a strong impact on toxicity of these elements, making them differentially inhibitory of nitrate-reducing versus aerobic cultures (Supplementary Dataset, Table S6). Several inorganic substrates of nitrate reductase are more inhibitory of aerobic cultures including AsO_4_^2−^, AsO_3_^2−^, CrO_4_^2−^, SeO_3_^2−^ and TeO_3_^2−^. Reduction of these compounds to less toxic products by *Pseudomonas* and other bacteria is known (Oremland and Stolz, 2000; Garbisu *et al.*, 1996). In contrast, TeO_4_^2−^, IO_3_^−^, IO_4_^−^ and BrO_3_^−^ are more inhibitory of nitrate-reducing cultures likely because they are reduced to more toxic compounds by nitrate-reductases. Because, Se, Te, Cr and As oxyanions can vary widely in their inhibitory potency depending on redox state of the element and whether N2E2 is respiring oxygen or nitrate, we suggest that these compounds should also be considered alongside molybdate limitation (Thorgersen *et al.*, 2015b) in future studies as possible constraints on growth and nitrate reduction in the most contaminated wells.

Co^2+^ is more inhibitory of aerobic cultures and Zn^2+^ is more inhibitory of nitrate-reducing cultures. The mechanisms of metal toxicity can be complex, but aerobically redox stress and the formation of reactive oxygen species is important (Lemire *et al.*, 2013). Co^2^+ is a better catalyst of Fenton chemistry than Zn^2+^ (Strlič *et al.*, 2003), while under nitrate-reducing conditions, Zn may interfere with Cu uptake and cofactor formation in the nitrite reductase NirK, or the nitrous oxide reductase Nos (Kraft *et al.*, 2011).

#### Iron and trace mineral depletion

Nutritional metals supply essential metallocofactors, and trace metal limitation can have a dramatic effect on microbial activity (Lemire *et al.*, 2013; Braud *et al.*, 2009). Thus, we measured the inhibitory potency of the 80 inorganic ions in an iron-limited rich media widely used in *Pseudomonas* studies, King’s B broth (KB) (Duffy and Défago, 1999) (Figure S4E) and in a minimal medium depleted in trace minerals (Figure S2F). Co^2^+, Sr^2^+, and VO_4_^2−^ are more inhibitory of N2E2 growth in KB versus LB and are known to interfere with iron uptake mechanisms (Figure S4) (Braud *et al.*, 2009). Ga^3^+, another iron analog (Lemire *et al.*, 2013) was slightly more inhibitory of KB cultures. The carbon sources in LB are simple sugars, amino acids and peptides whereas in KB they are glycerol and amino acids. As with our comparison of lactate and glucose as a carbon source, Be^2^+ is more inhibitory of cultures relying on substrate-level phosphorylation (e.g. LB, glucose defined) (Klemperer, 1950). VO_4_^2−^ is more inhibitory of KB cultures and in this context it may be both an iron analog (Wichard *et al.*, 2009; Braud *et al.*, 2009) as well as a phosphate analog (Benabe *et al.*, 1987).

Several compounds are more inhibitory of trace mineral depleted cultures (Figure S2F). Ni^2^+, Al^3^+, and Pb^2^+ interfere with transition metal uptake (Teitzel and Parsek, 2003; Braud *et al.*, 2009) while TeO_4_^2−^ and WO_4_^2−^ can interfere with molybdate or sulfate uptake both of which are depleted in these conditions (Wichard *et al.*, 2009; Enoch and Lester, 1972; Prins *et al.*, 1980). CN^−^ is nearly 3 orders of magnitude more inhibitory of mineral replete cultures. CN^−^ binds to respiratory metallocofactors, but its degradation is catalyzed by metal siderophore complexes under metal depleted conditions by *Pseudomonas* (Luque-Almagro *et al.*, 2005; Chen and Kunz, 1997).

Further work will evaluate how trace nutrient depletion impacts microbial communities at the ORFRC. For example, molybdenum limitation may limit nitrate reduction activity in some areas of the ORFRC and select for organisms with higher molydenum affinity (Thorgersen *et al.*, 2015a). By extending the approach we used in this study, a high-throughput dose-response strategy can be used to both determine nutrient limitation thresholds for nitrogen, phosphate, sulfur and trace vitamins/minerals and systematically determine toxicity thresholds for specific inhibitors of nutrient uptake systems (Carlson *et al.*, 2017).

## Supplementary Figures

**Figure S1.**
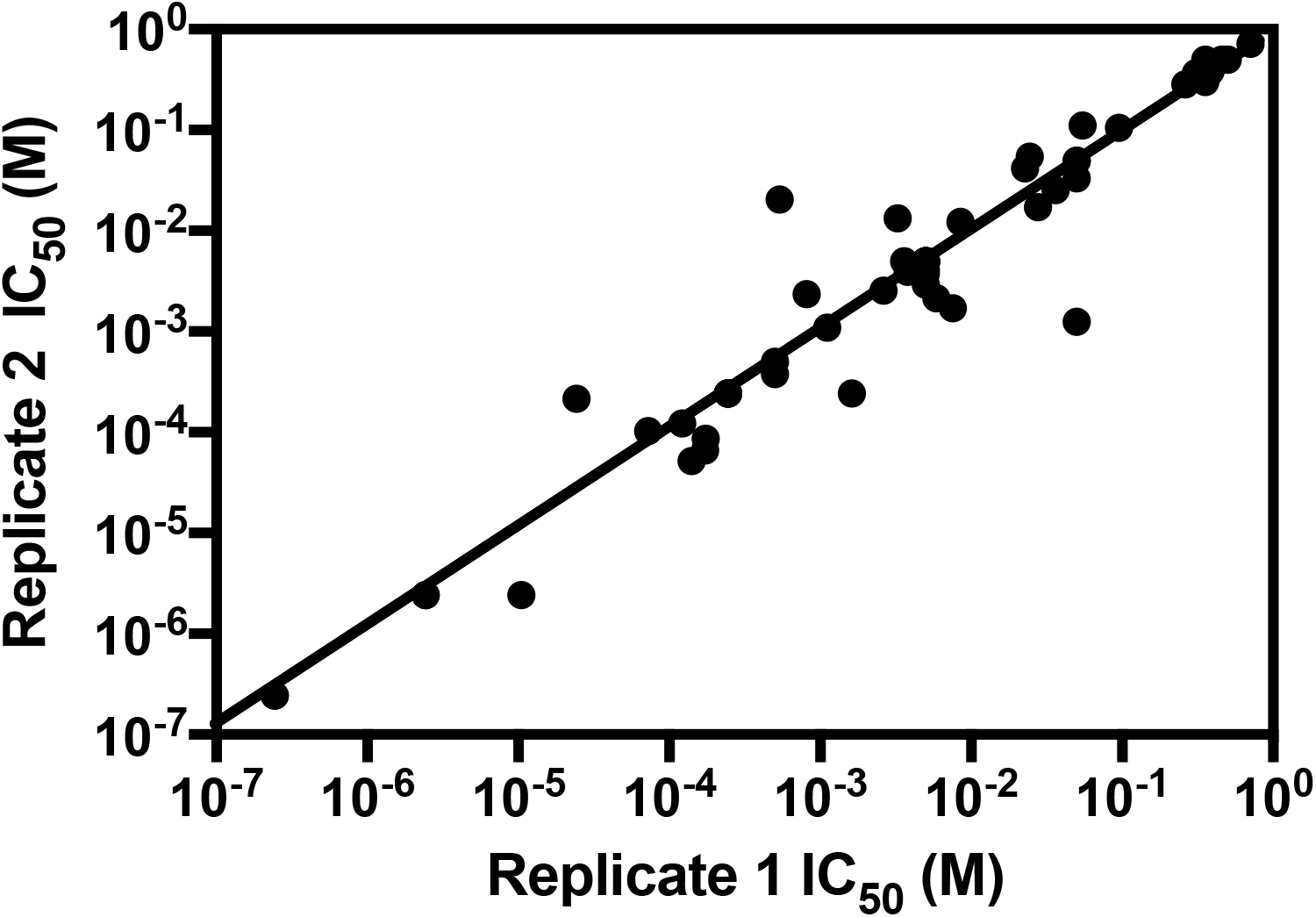
IC_50_s measured in replicate rich media (LB/aerobic) dose-response assays conducted on different days. All compounds have overlapping 95% confidence intervals and are not selective (Materials and Methods).

**Figure S2.**
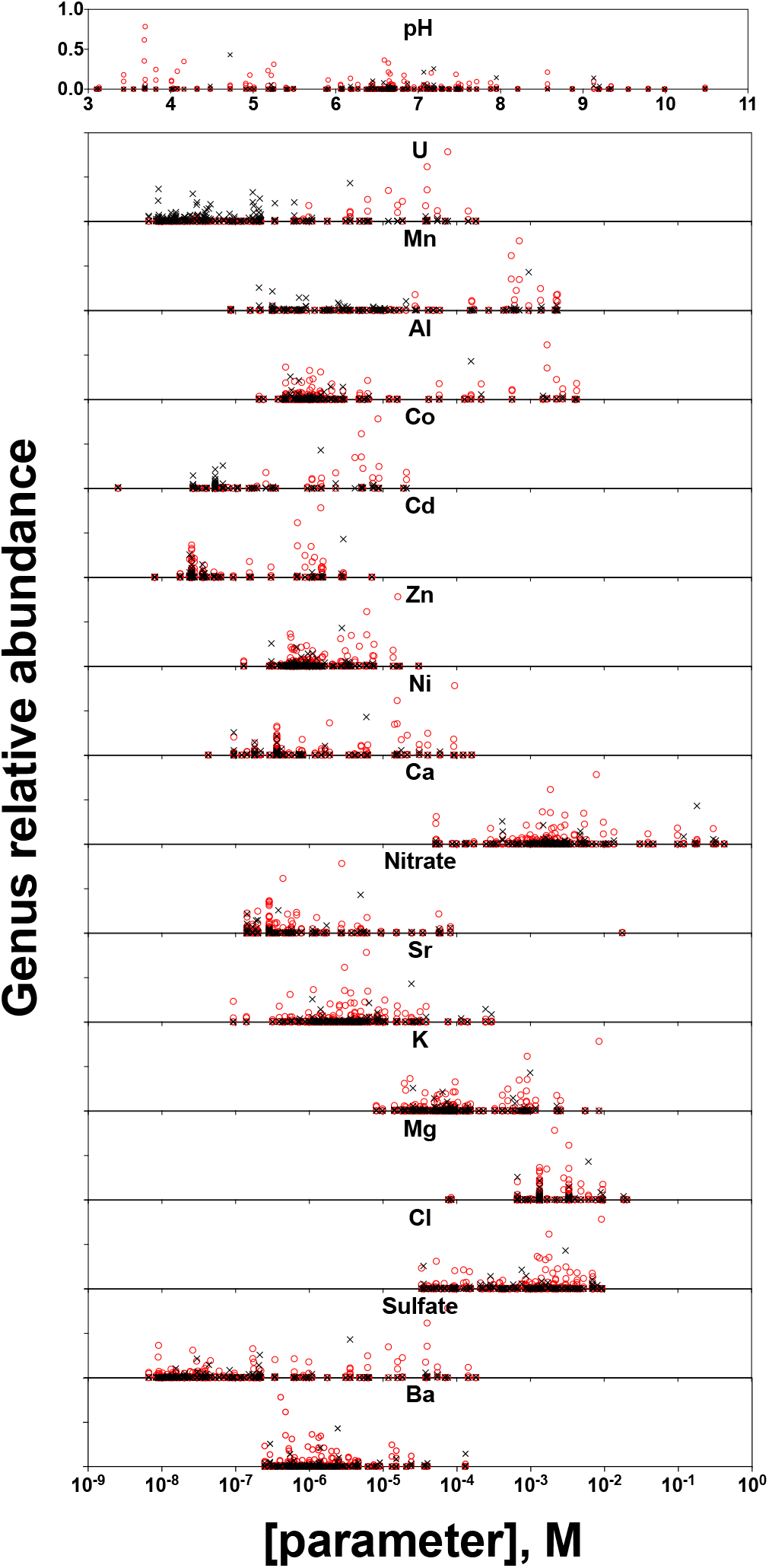
Relative abundances of *Rhodanobacter* (red circles) and *Pseudomonas* (black x’s) plotted versus concentrations of selected geochemical parameters significantly correlated with high *Rhodanobacter* relative abundance (Spearman’s rho, adjusted p-value >0.05).

**Figure S3.**
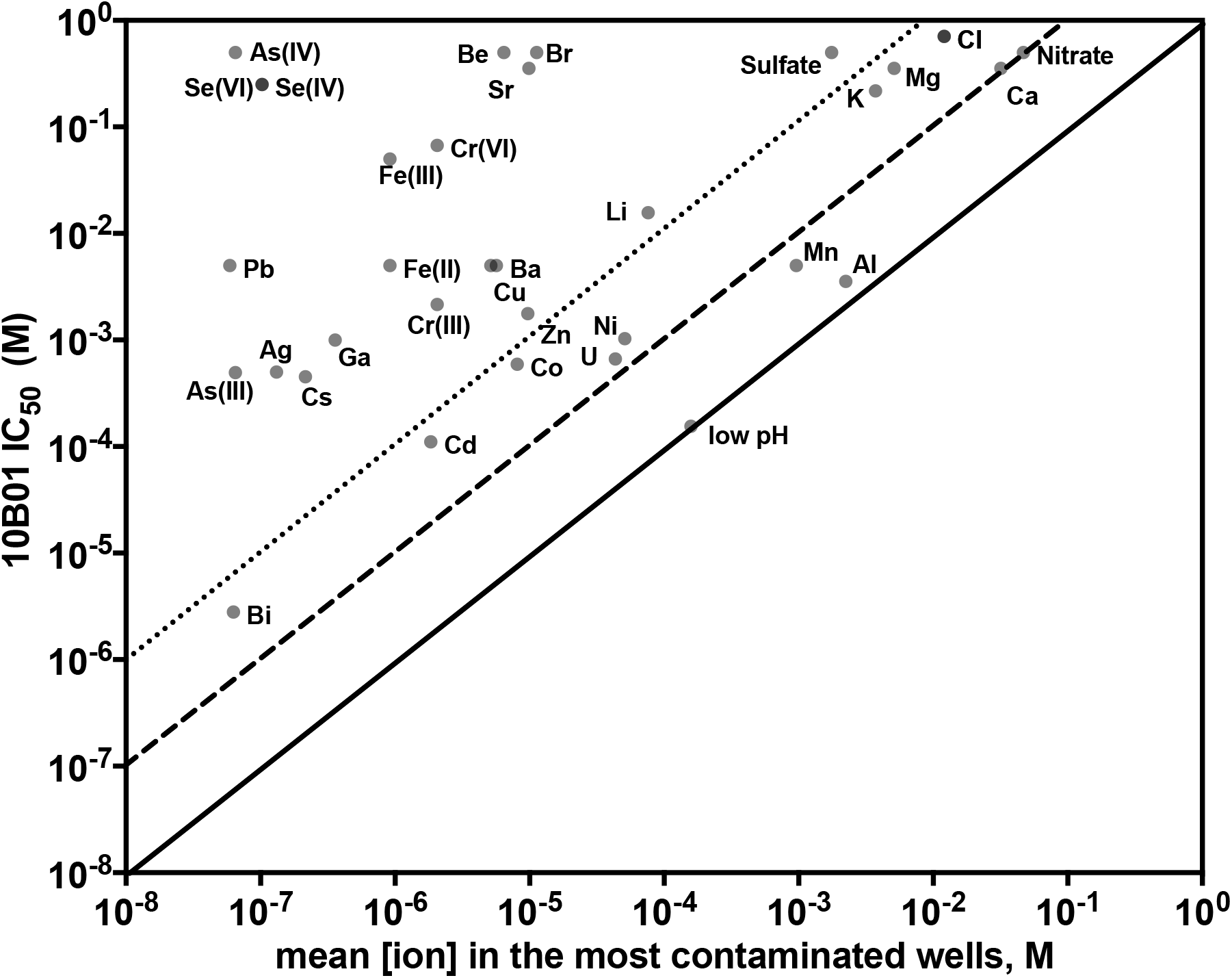
*Rhodanobacter sp.* FW104-10B01 (10B01) IC_50_s compared with average concentrations of inorganic ions in groundwater samples with >5% *Rhodanobacter*. pH is represented by [H_3_O^+^].

**Figure S4.**
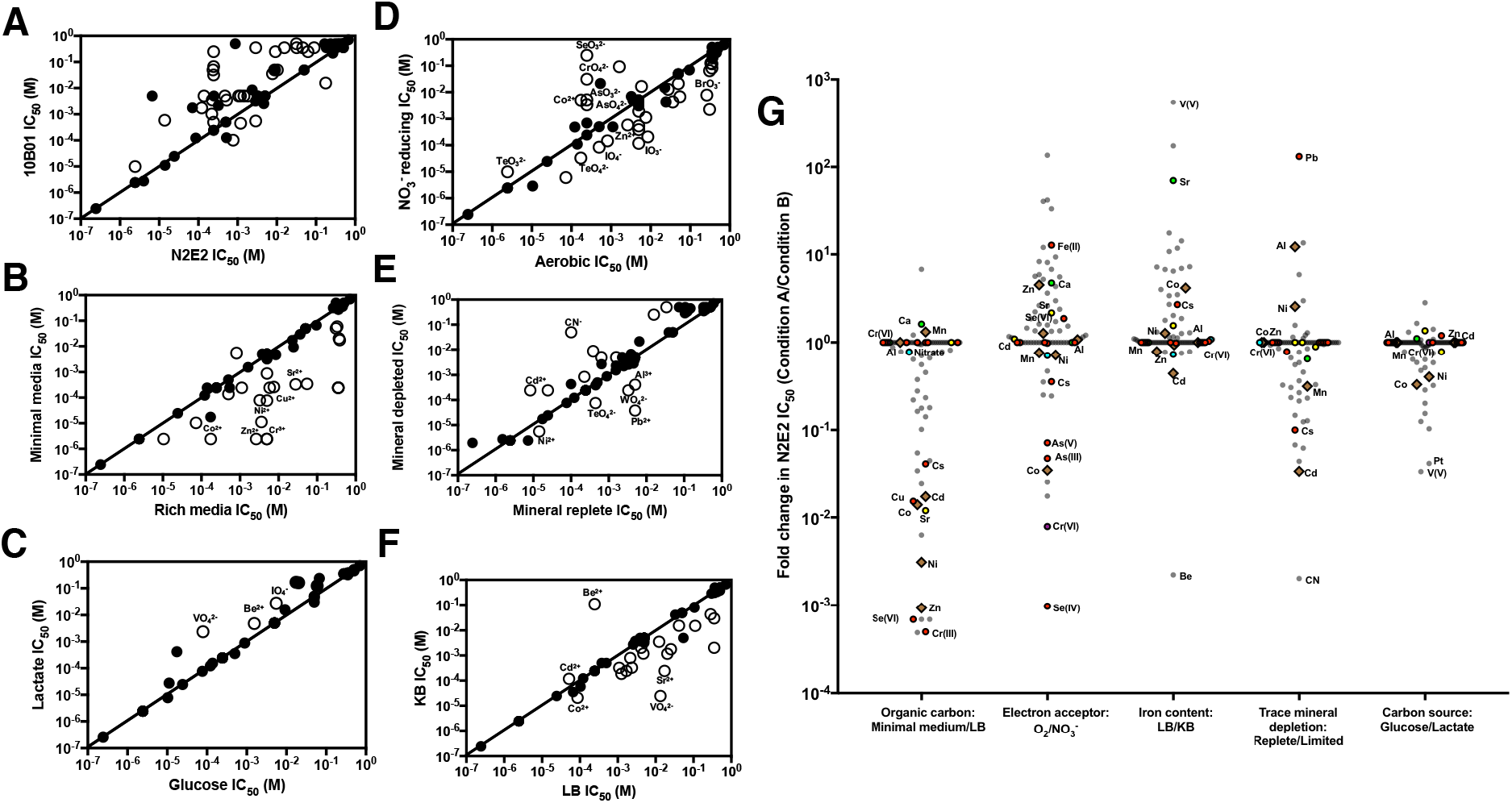
Influence of growth medium conditions on compound inhibitory potencies against *Pseudomonas fluorescens* FW300-N2E2 (N2E2). Selected compounds with altered inhibitory potency in various growth condition comparisons are labelled. **A**. IC_50_s for compounds against N2E2 and 10B01 grown in R2A medium. **B**. IC_50_s for compounds against N2E2 grown in a chemically defined minimal medium or rich medium (Luria Broth, LB). **C**. IC_50_s for compounds against N2E2 grown in minimal media with lactate or glucose as a carbon source. **D**. IC_50_s for compounds against N2E2 grown in LB aerobically or anaerobically with nitrate as the sole electron acceptor. **E**. IC_50_s for compounds against N2E2 grown in minimal media in mineral depleted conditions (0.25x trace mineral stock) or mineral replete conditions. **F**. IC_50_s for compounds against N2E2 grown in King’s B media (KB) or LB. **G**. The fold change in N2E2 inorganic IC_50_s is plotted for various growth media conditions. Coloring and symbols is as in Figure 3C.

**Figure S5.**
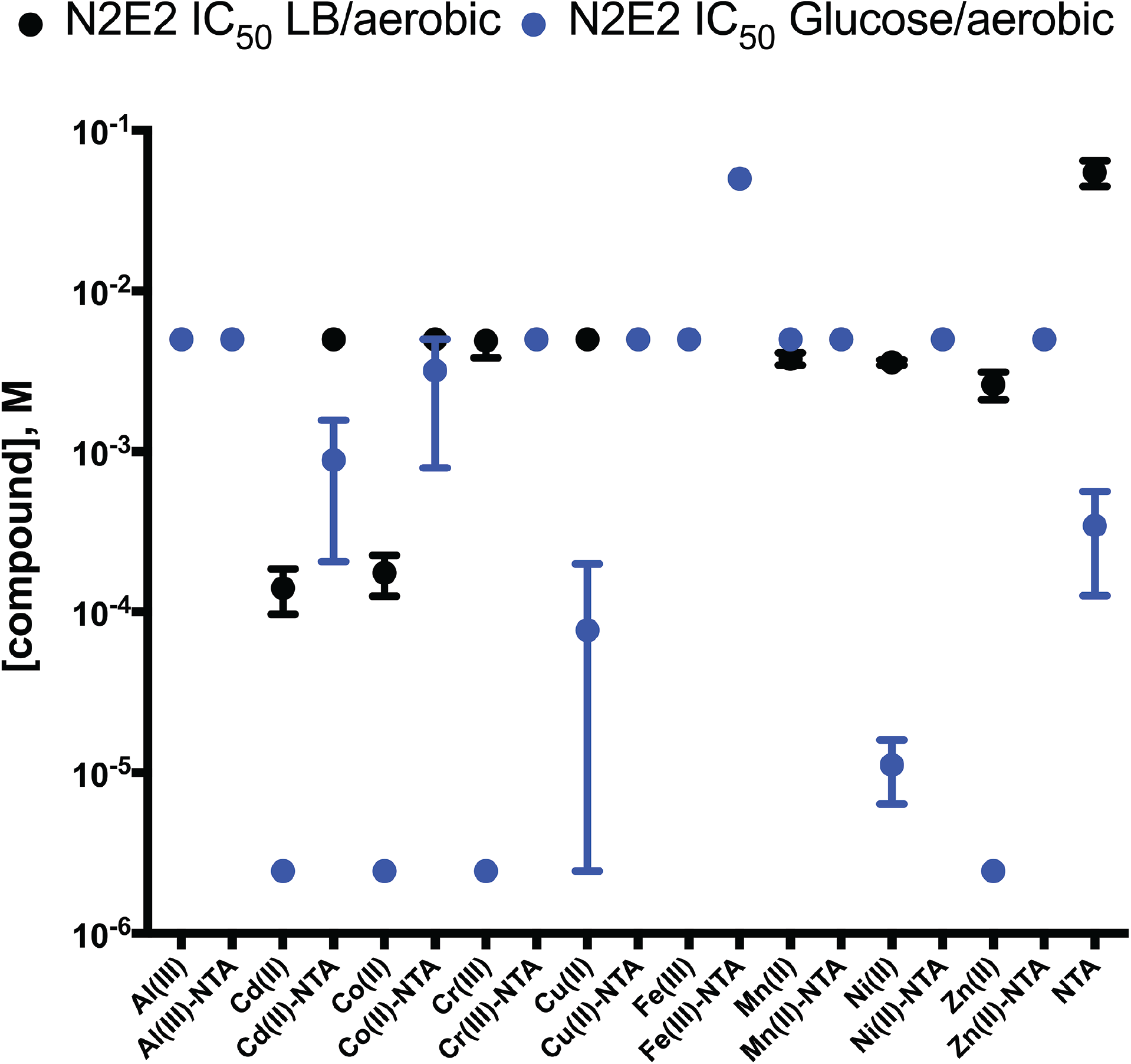
Influence of chelation on transition metal inhibitory potency. Inhibitory potencies (IC_50_, 95% CI) of selected free ions or NTA complexes against *Pseudomonas fluorescens* FW300-N2E2 grown aerobically in either rich media (LB/aerobic) or minimal media (Glucose/aerobic).

**Figure S6.**
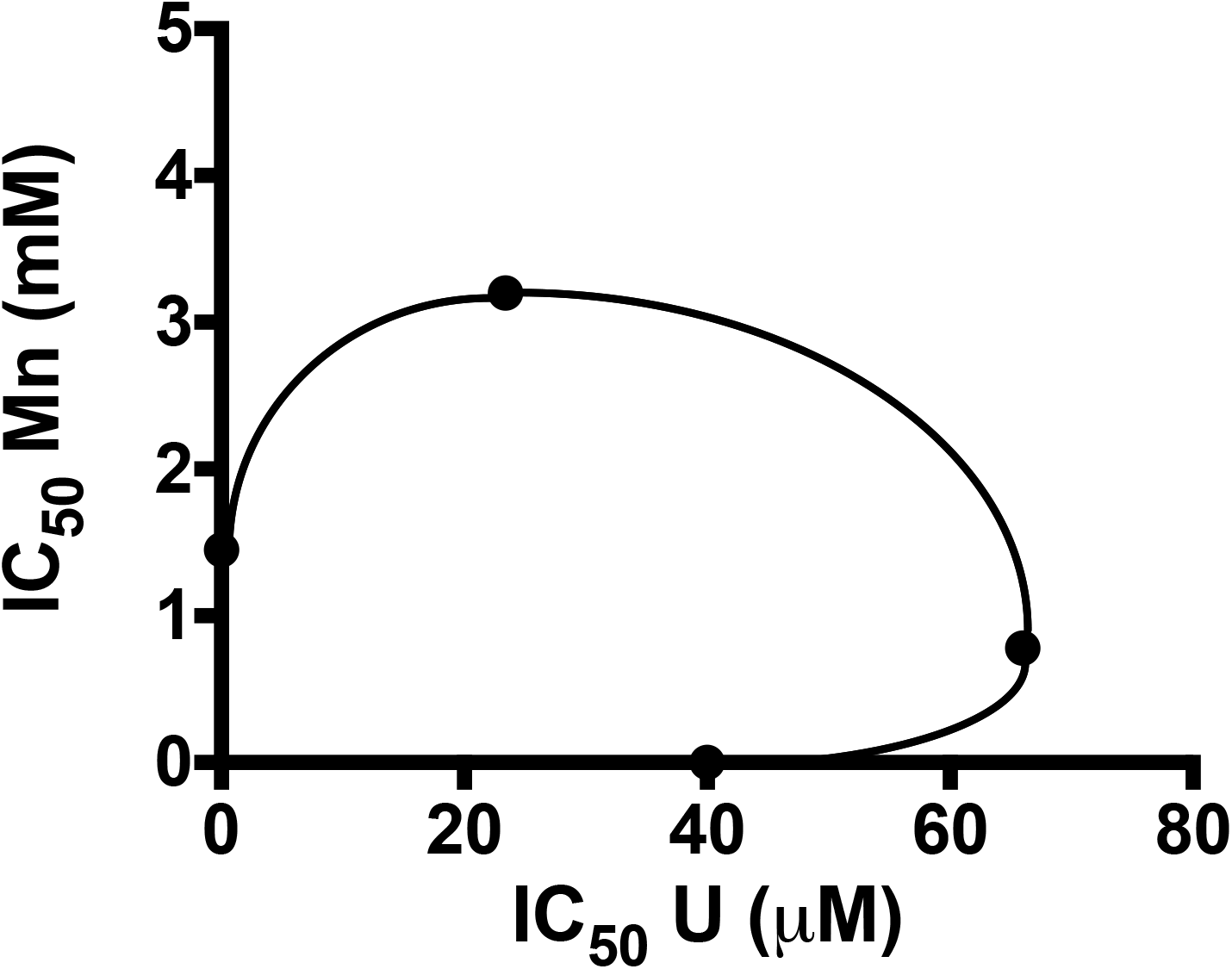
Antagonistic inhibition of *Pseudomonas fluorescens* FW300-N2E2 (N2E2) for combinations of Mn and U. The fractional inhibition concentration index (FICI, Supplementary Materials and Methods) is reported.

## Notes

**Grant information:** This work was funded by ENIGMA, a Scientific Focus Area Program supported by the U. S. Department of Energy, Office of Science, Office of Biological and Environmental Research, Genomics: GTL Foundational Science through contract DE-AC02-05CH11231 between Lawrence Berkeley National Laboratory and the U. S. Department of Energy.

